# Top-down control of sustained attention by the medial prefrontal cortex (mPFC) - locus coeruleus (LC) circuit during the rodent continuous performance test (rCPT)

**DOI:** 10.64898/2025.12.01.691673

**Authors:** Jason J. Rehg, Daniel E. Olivares, Ye Li, Keri Martinowich, Gregory V. Carr, Jorge Miranda-Barrientos

## Abstract

The medial prefrontal cortex (mPFC) plays a pivotal role in attention by exerting top-down control to allocate cognitive resources toward behaviorally relevant stimuli based on learned context and expectations. mPFC neurons project to multiple cortical and subcortical regions, including the locus coeruleus (LC)—the brain’s primary source of norepinephrine (NE). The mPFC also receives inputs from the LC, which release NE to modulate mPFC neuronal activity and downstream cellular signaling. While enhanced functional connectivity between the mPFC and LC in mice during sustained attention tasks suggest an important role for the mPFC–LC circuit, functional evidence directly implicating this circuit in attention is lacking. Here, we investigated the role of the mPFC–LC circuit in attention by comparing selective chemogenetic manipulation of mPFC neurons that project to the LC (mPFC-LC projectors) to non-specific chemogenetic manipulation of mPFC neurons. Selective activation of mPFC–LC projectors in mice performing the rodent continuous performance test (rCPT), a translational sustained attention task, robustly improves attentional performance by enhancing discrimination while non-selective activation of mPFC neurons increases attentional performance by increasing responsiveness. Behavioral effects of mPFC-LC projector activation were mediated by recruitment of a microcircuit involving LC-NE neurons and glutamate and GABA peri-LC neurons while effects of non-selective activation of mPFC neurons were mediated by engaging downstream targets such as the nucleus accumbens (NAc) as well as the LC/peri-LC region.

## Introduction

The medial prefrontal cortex (mPFC) plays a pivotal role in numerous cognitive functions, including attention, decision-making, learning, and memory [1–3]. The mPFC exerts top-down attentional control by sending descending feedback signals that bias sensory processing toward behaviorally-relevant information, guided by learned context, expectations, and goal-directed selection [4–7]. mPFC dysfunction is implicated in multiple neuropsychiatric disorders, including attention-deficit/hyperactivity disorder (ADHD) and schizophrenia [3,8], both of which are characterized by impaired attentional control [9,10].

The mPFC sends projections to numerous cortical and subcortical targets, forming circuits that control attention [9,11–13]. Among these, the mPFC sends glutamatergic axons to the locus coeruleus (LC)—the brain’s primary source of norepinephrine (NE)—which synapses onto LC-NE neurons [14,15]. Additionally, mPFC axon terminals contact GABAergic neurons in the peri-LC region, which in turn modulate LC-NE activity, forming a complex regulatory system [15,16]. LC-NE neurons project back to the mPFC, where downstream NE signaling regulates multiple facets of attention, including arousal [17,18], cognitive flexibility [19], adaptive gain [20], and inhibitory control [21]. Despite the well-established role of NE in attentional processes, the specific contribution and mechanisms by which mPFC projections to the LC regulate attentional control remain unclear. Recently, we showed that mPFC–LC oscillatory activity was associated with performance during the rodent continuous performance test (rCPT) [22]—a well-validated translational paradigm of sustained attention in which animals must discriminate between target and non-target stimuli across successive trials [23–25]. While this correlation suggests an important role for the mPFC–LC circuit in attentional control, functional evidence that establishes causality of this circuit is still lacking.

Here, we investigated the role of the mPFC–LC circuit in attentional control by chemogenetically manipulating the activity of mPFC-LC projectors in mice performing the rCPT. This data was compared to mice performing the rCPT after broad manipulation of mPFC neuronal activity. Selective activation of mPFC–LC projectors robustly improves attentional performance by enhancing discrimination without affecting responsiveness or impulsivity. In contrast, general activation of mPFC neurons alters multiple aspects of attention, including responsiveness, discrimination, and impulsivity. Furthermore, we found that mPFC–LC projectors exert their effects by directly activating LC-NE neurons and differentially engaging excitatory and inhibitory peri-LC cell types, forming a complex regulatory microcircuit. In contrast, broad activation of mPFC neurons appears to mediate its behavioral effects through a wider set of downstream circuits that includes not only the LC, but also regions in the mesolimbic reward system related to motivational drive such as the nucleus accumbens (NAc).

## Materials and Methods

### Animals

Male C57BL/6J mice (Strain #000664; The Jackson Laboratory, Bar Harbor, ME) were 8-10 weeks old at the start of experiments. Mice were group housed (4/cage) in disposable polycarbonate caging (Innovive, San Diego, CA) and maintained on a reverse 12/12 light/dark cycle (lights on at 19:00 hours, lights off at 07:00 hours). For behavior experiments, mice were housed 2/cage following surgeries for virus injection to minimize competition for food during restriction. Mice received Teklad Irradiated Global 16% Protein Rodent Diet (#2916; Envigo, Indianapolis, IN) in the home cage ad libitum until the start of the food restriction protocol, and water was available in the home cage ad libitum throughout all experiments. Behavioral testing was conducted Monday-Friday during the dark phase (07:00-19:00 hours). All experiments and procedures were approved by the Johns Hopkins Animal Care and Use Committee and in accordance with the Guide for the Care and Use of Laboratory Animals.

### Surgical procedures

For viral injection surgeries, mice were anesthetized with isoflurane (induction: 2-4% in oxygen, maintenance: 1-2%) and secured to a stereotaxic frame. The top of the skull was exposed by an incision along the midline of the scalp, Bregma and Lambda were identified, and the head was leveled to ensure the skull was flat. A small hole was drilled with a 0.9 mm burr (Fine Science Tools, Foster City, CA) above each injection site (**mPFC:** AP: +1.8, ML: ±0.3, DV: -1.7; **LC:** AP: -5.4, ML: ±0.9, DV: -3.5), and 400 nl of viral vector was injected. For general mPFC manipulations, a viral vector expressing excitatory (AAV5-hsyn-hM3Dq; Addgene) or inhibitory (AAV5-hsyn-hM4Di; Addgene) Designer Receptors Exclusively Activated by Designer Drugs (DREADDs) or reporter fluorophore (AAV5-hsyn-mCherry; addgene) was injected bilaterally in the mPFC. For mPFC-LC projector manipulations, a retrograde viral vector containing the recombinase *Cre* (AAVrg-hSyn-Cre; Addgene) was injected bilaterally at the LC, and a *Cre-*dependent virus containing excitatory (AAV5-hsyn-DIO-hM3Dq; Addgene), inhibitory (AAV5-hsyn-DIO-hM4Di; Addgene) DREADDs or an mCherry reporter (AAV5-hSyn-DIO-mCherry) was injected bilaterally in the mPFC. Injections were made using a Micro4 controller and UltraMicroPump along with a 10 µl Nanofil syringes equipped with 33-gauge needles (WPI Inc., Sarasota, FL). The syringe was left in place for 10 min after injection to minimize diffusion. Following surgery, the incision site was closed using surgical staples (Fine Science Tools, Foster City, CA), and animals recovered on a heating pad for 20-40 min before being returned to the colony room. Animals were closely monitored for health and recovery progress and received Meloxicam injections (20 mg/kg) to relieve pain for three additional days.

### Food restriction protocol

Before initiating behavior training, mice were placed on a food restriction protocol to increase motivation in the task and avoid satiety-related issues during performance. Mice were handled and weighed for at least two consecutive days to establish a baseline weight, then food restricted to 2.5-3 g of chow per mouse per day and weighed daily to ensure that they maintained at least 85-90% of their free-feeding weight based on baseline weight and average growth curve data for the strain (The Jackson Laboratory, Bar Harbor, ME). To familiarize the mice to Nesquik® strawberry milk (Nestlé, Vevey, Switzerland), which was used as reward during rCPT training, a 4×4 cm weighing plate (VWR, Radnor, PA, USA) containing ∼2 ml of strawberry milk was introduced to the home cage for three consecutive days. The weighing plate was left in the cage until all mice had sampled the strawberry milk.

### Behavior training

Behavior training started at least one week after surgeries to allow recovery and minimize stress to the animals. After the recovery period, mice were subjected to 3 days of handling sessions consisting of 5 min sessions each day to accustom the mice to the experimenter.

#### Habituation

After handling sessions, mice were given two consecutive habituation sessions (20 min each) to familiarize them to the Bussey-Saksida mouse touchscreen chambers (Lafayette Instruments, Lafayette, IN). In habituation sessions, 1 mL of strawberry milk was placed into the reward tray. The screen was responsive to touch, but touches were not rewarded.

#### Rodent Continuous Performance Test (rCPT)

rCPT training protocol was based on a previously described protocol [[26]]. Briefly, mice were trained in the touchscreen chambers, which were connected to a computer running ABET II software (Campden Instruments, Loughborough, UK) to track behavioral responses during rCPT sessions.

##### Stage 1

Mice received 45 min training sessions (Monday-Friday), during which they learned to respond to a visual stimulus (white square) presented at the center of the touch screen. The stimulus was displayed for 10 s (stimulus duration, SD), during which a touch in the center of the screen produced delivery of ∼20 ul of strawberry milk in the reward tray located on the opposite side of the chamber. Following SD, a 0.5 s limited hold (LH) period was given in which the screen was blank, but a touch would still yield a reward. Upon interacting with the stimulus, a 1 s tone (3 kHz) was delivered, the reward tray was illuminated signaling reward delivery, and the schedule was paused until a head entry into the reward tray was detected by an IR beam. Then, a 2 s intertrial interval (ITI) would begin before the subsequent trial started. If the mouse did not interact with the stimulus during the SD or LH, an ITI would start, and the next trial would follow. The criterion for a mouse to advance to the next stage was to obtain at least 60 rewards per session in two consecutive sessions.

##### Stage 2

In Stage 2, a target stimulus (S+) was introduced. The S+ consisted of a square with either horizontal or vertical black and white bars that replaced the white square at the center of the screen. Sessions were 45 min long, and each mouse was assigned either horizontal or vertical oriented S+ for the remaining sessions of the experiment. The S+ assignment was counterbalanced. During Stage 2, the SD was reduced to 2 s, and LH was increased to 2.5 s. Interaction with S+ (hit) during SD + LH resulted in reward delivery. Once a mouse obtained at least 60 hits / session in two consecutive sessions, it was moved to the next stage.

##### Stage 3

In Stage 3, a non-target stimulus (S-) consisting of a snowflake shape presented at the center of the screen was introduced. On each trial, the probability of S+ / S- was 50% / 50%. The SD and LH were identical to Stage 2, but the ITI length was either 2 or 3 s in length (randomized ITI duration during trials). Similar to Stage 2, screen touches during S+ (hit) yielded a reward but not screen touches during S- (false alarm (FA)). A FA resulted in the beginning of the ITI followed by a correction trial. In correction trials, a S- was presented again. If another FA occurs, a new correction trial starts until the mouse doesn’t interact with the S- (correct-rejection). We used discrimination index (d’) which is a measure of sensitivity bias (refers to the perceptual discriminability between the S+ and S−) to determine attention performance during Stage 3. Mice were trained in Stage 3 until they reached stable performance, as defined by a d’ score of 0.6 or higher over 3 consecutive sessions.

### Experimental Probes

#### S3 probe

Once mice reached stable performance in Stage 3, they progressed to the Stage 3 probe (S3 probe), in which the effect of chemogenetic manipulations on performance was assessed using the same task parameters on which mice trained in S3. Mice completed 4 consecutive days of 45 min testing sessions, receiving either saline or J60 injections intraperitoneally 15 min before the session on alternating days (Saline M/W; J60 T/Th). Two testing sessions were performed for each treatment to account for day-to-day fluctuations in performance and metrics were averaged between same-treatment sessions.

#### Degraded Stimulus Probe

The degraded stimulus probe was carried out over two weeks of testing immediately following the S3 probe. Each week consisted of 4 days of testing, alternating between sessions with the regular stimulus (M/W) and probe sessions with the degraded stimulus (T/Th). Mice were randomly assigned to receive J60 before degraded stimulus sessions on either the first week or the second week, and received saline injections before all other sessions.

#### *Time-on-Task* (TOT) *Probe*

For the time-on-task probe, mice underwent an additional week of testing in which they alternated between standard 45 min sessions (M/W) and extended 90 min sessions (T/Th). All mice were randomly assigned to receive J60 before either the first or the second extended session, receiving saline injections prior to all other testing sessions. For extended sessions, performance data was subdivided into 15 min time bins for analysis.

### DREADDs activation for *Fos* quantification

To evaluate *Fos* expression following DREADDs activation, mice were injected with JHU37160 (J60, 0.5 mg/kg i.p.), 15 min prior to being placed into touchscreen chambers for a 45 min S3 rCPT session. 20 min after completion of the rCPT session, mice were sacrificed by cervical dislocation and brains were then extracted. Tissue was flash frozen in 2-methylbutane (ThermoFisher) and stored at -80°C until cryosectioning.

### Multiplexed Fluorescent in situ Hybridization (RNAscope)

Coronal sections (12 μm) were collected across the rostral-caudal axis of the mPFC and mounted onto slides (VWR, SuperFrost Plus) to verify viral expression and in the LC/peri-LC region, ventral tegmental area (VTA) and NAc for analysis of cell types activated by mPFC-LC projectors or broad mPFC neurons. To identify activated cell types, we used the RNAScope Fluorescent Multiplex Kit V2 (Advanced Cell Diagnostics[ACD]). The slides were fixed in 10% buffered formalin at 4°C, washed in 1x PBS, and dehydrated in ethanol. Slides were pre-treated with protease IV solution, and subsequently incubated at 40°C for 2 hours in a HybEZ oven (ACD) with antisense probes against *Th* (Cat No. 317621), *Slc17a6* (Cat No. 319171), *Slc32a1* (Cat No. 319191), and *Fos* (Cat No. 316921). Transcript expression was visualized using AXNikonTi2-Econfocal fluorescence microscope equipped with NIS-Elements(v5.42.02). For cell type quantification, a 20X tiled, z-stacked image of the entire LC region from each mouse was acquired at atlas-matched coordinates along the rostrocaudal axis (see **Fig. S5**), and images were saved as ND2 files containing unstitched z-stacks.

### Quantitative and Statistical Methods

#### Behavioral Scoring

Behavioral databases containing the timestamps from stimuli presentation, hits, false alarms, latency to response, etc. were retrieved from ABET II (Lafayette Instruments, Lafayette, IN) and Whisker server (Cambridge University Technical Services, UK). Behavioral data was analyzed using Excel to obtain performance scoring parameters. Performance scoring parameters were similar to those described in [25] and [27]. Briefly, to assess attention performance during Stage 3 training and experimental probes, we calculated discrimination index ***d’*** and response criterion ***c*** with the following formulas:

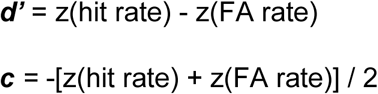

Whereas:

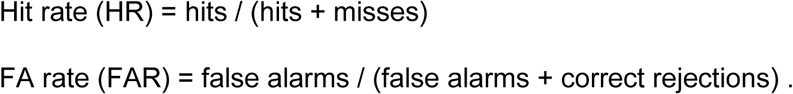

#### Statistical analysis

All statistical analyses were performed in Prism v10.2.0 (GraphPad Software, Inc.) or RStudio (2025.05.0+496). Behavioral metrics were analyzed within-subject using paired *t*-tests for session totals and 2-way analysis of variance (ANOVA) for time-binned data to evaluate mixed effects of treatment and time. FISH data was tested for normality using the Shapiro-Wilkes test, then analyzed using unpaired *t*-tests or ANOVA, where appropriate. All analyses were two-tailed and p values were adjusted for multiple comparisons where appropriate. Values are expressed as the means ± S.E.M. Statistical significance was accepted when p<0.05.

## Results

### Activation of mPFC-LC projectors improves performance in the rCPT by increasing discrimination

We recently demonstrated that oscillatory activity across the mPFC-LC circuit is associated with attentional behavior in the rCPT [28]. To test the functional role of this circuit in sustained attention, we chemogenetically manipulated activity of mPFC-LC projectors in mice performing the rCPT by selectively expressing excitatory (hM3Dq) or inhibitory (hM4Di) DREADDs in mPFC-LC projectors. Specifically, we bilaterally injected C57BL/6J mice with a retrograde viral vector encoding Cre into the LC and a Cre-dependent viral vector encoding either hM3D(Gq)-mCherry (mPFC-LC^hM3Dq^), hM4D(Gi)-mCherry (mPFC-LC^hM4Di^), or mCherry alone (mPFC-LC^mCherry^) into the mPFC (**Fig. 1A**). Mice were trained in the rCPT until they achieved stable, proficient performance (d’ ≥ 0.6) on Stage 3 (S3). All groups successfully acquired the task with no significant differences in performance (**Fig. S1**). To assess the effect of manipulating activity of mPFC-LC projectors on rCPT performance, we conducted a 4 day probe during which mice received alternating injections of the DREADD ligand JHU37160 (J60; 0.5 mg/kg, i.p.) or saline during S3 sessions (S3-probe; **Fig. 1D**). During J60 sessions, DREADDs activation in mPFC-LC^hM3Dq^ group significantly increased d’ (*SAL*=2.05±0.10, *J60*=2.51±0.10; t(11)=4.383, p=0.0011; **Fig. 2C**), driven by a reduction in the false alarm rate (FAR; *SAL*=0.12±0.02, *J60*=0.08±0.01; t(11)=3.111, p=0.0099), while hit rate (HR) remained unchanged (*SAL*=0.77±0.03, *J60*=0.81±0.03; t(11)=1.175, p=0.2648). Notably, J60 administration had no significant effect on mean total responses (*SAL*=176.25±7.22, *J60*=178.15±6.96; t(11)=1.073, p=0.3063), ITI touches (*SAL*=55.35±10.79, *J60*=59.20±16.04; t(11)=0.3852, p=0.7090), or response criterion (*c*-parameter) (*SAL*=0.23±0.09, *J60*=0.24±0.09; t(11)=0.8658, p=0.4051). Together, these findings indicate that chemogenetic activation of mPFC-LC projectors enhances rCPT performance by improving discrimination without affecting responsiveness or response strategy (**Fig. 2C**). Interestingly, J60 administration in mPFC-LC^hM4Di^ mice induced a strong trend towards increased d’ (*SAL*=1.84±0.13, *J60*=2.09±0.14; t(11)=2.050, p=0.0649), which was driven by an increased HR (*SAL*=0.67±0.05, *J60*=0.73±0.04; t(11)=2.005, p=0.0702) (**Fig. 2D**). Similar to mPFC-LC^hM3Dq^ mice, J60 administration did not affect any other performance metric in mPFC-LC^hM4Di^ mice (FAR: *SAL*=0.11±0.02, *J60*=0.09±0.01, t(11)=0.9483, p=0.3634; c: *SAL*=0.41±0.09, *J60*=0.35±0.08, t(11)=1.007, p=0.3355; responses: *SAL*=150.04±9.99, *J60*=160.50±9.06, t(11)=1.606, p=0.1366). mPFC-LC^mCherry^ group showed no effect on any rCPT performance metric between J60 and saline sessions (d’: *SAL*=1.81±0.12, *J60*=1.79±0.15; t(7) = 0.6531, p=0.5346; HR: *SAL*=0.65±0.04, *J60*=0.61±0.05; t(7)=1.230, p=0.2585; FAR: *SAL*=0.10±0.01, *J60*=0.09±0.01; t(7)=1.174, p=0.2790; c: *SAL*=0.22±0.09, *J60*=0.31±0.13; t(7)=1.477, p=0.1832; responses: *SAL*=177.31±9.36, *J60*=171.56±11.69; t(7)=1.272, p=0.2440)(**Fig. S2**).

**Figure 1.**
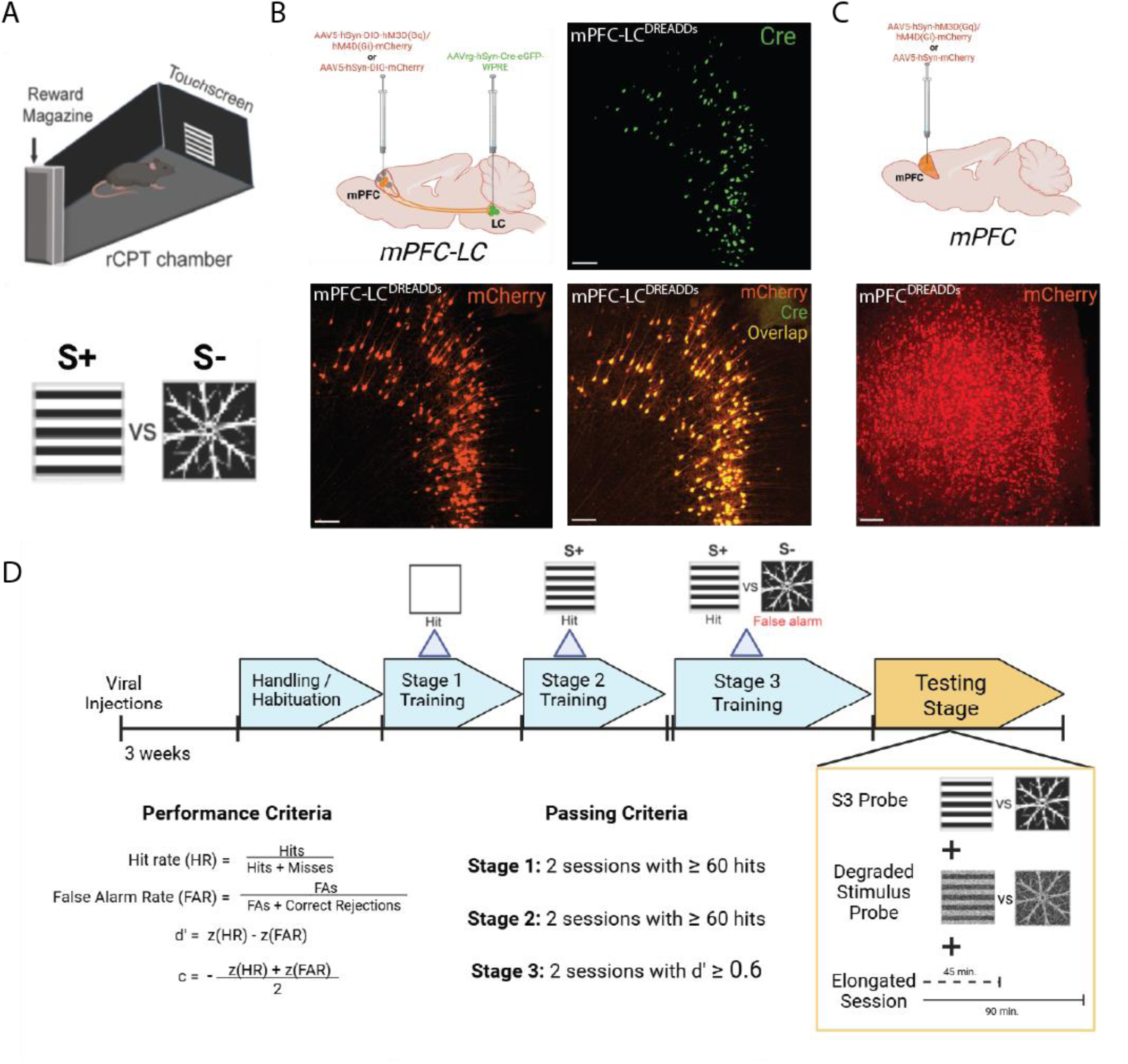
Study design and overview of the rCPT. **A.** Schematic of rCPT chamber along with target (S+) and non-target (S-) Stage 3 stimuli. **B.** Schematic of viral strategy to selectively express DREADDs in mPFC-LC projectors (top-left) and mPFC micrographs showing Cre-GFP (top-right), DREADDs-mCherry (bottom-left) and the merge image showing colocalization of Cre-GFP and DREADDs-mCherry in mPFC-LC projectors (bottom-right) in mPFC-LC projectors. Scale bars = 100µm. **C.** Schematic of viral strategy to express DREADDs pan-neuronally within the mPFC (top) and mPFC micrographs showing DREADDs-mCherry (bottom) in mPFC neurons. Scale bars = 100µm. **D.** Timeline for rCPT training (top) and schematic of testing probes (bottom right). Formulas for calculating performance during rCPT (bottom left), and passing criteria (bottom center) for each rCPT training phase.

**Figure 2.**
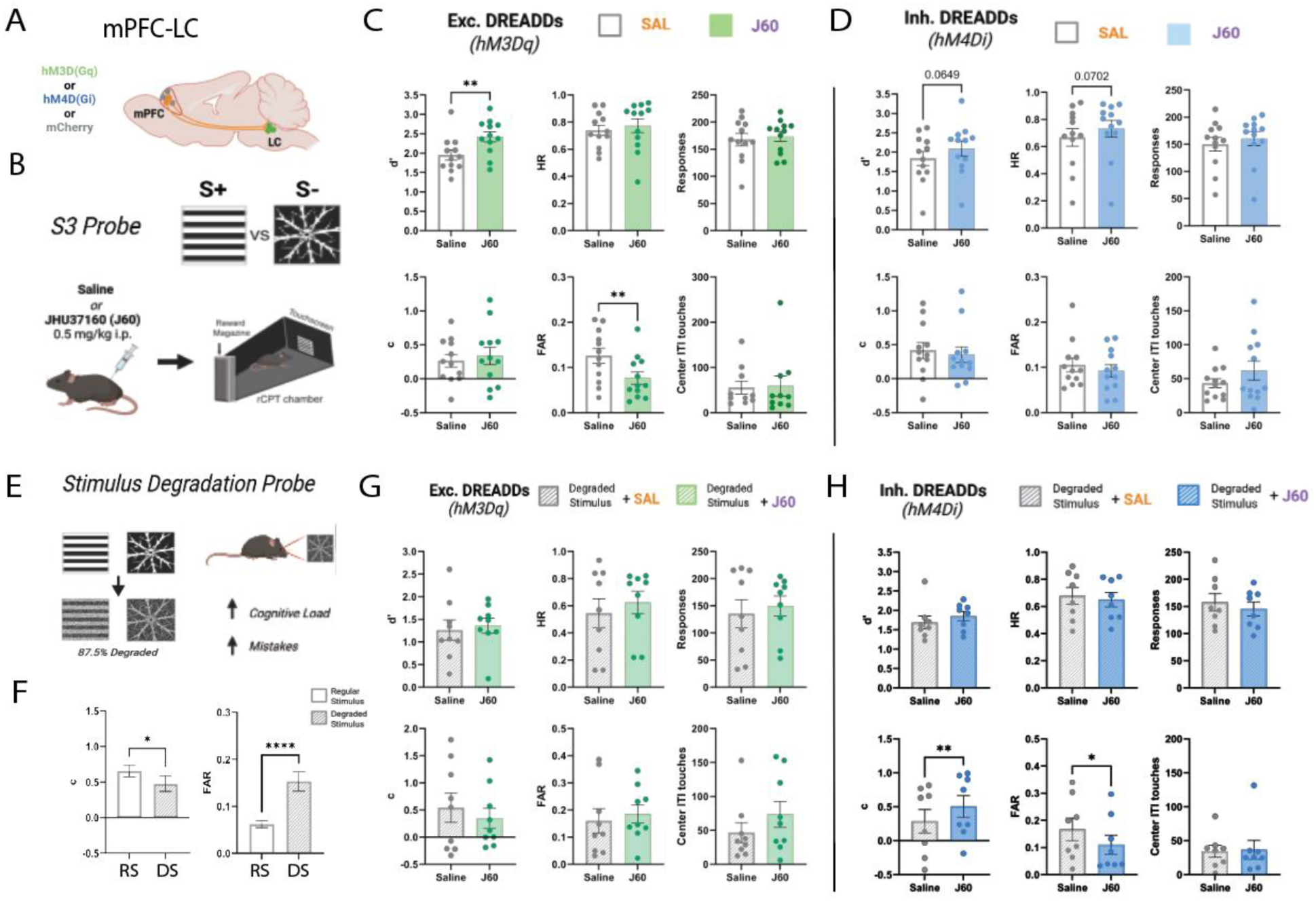
Selective activation of the mPFC-LC circuit enhances discrimination in the rCPT while inhibition improves performance when cognitive load is increased. **A.** Representative schematic of the mPFC-LC circuit. **B.** Overview of S3 probe with visual stimuli and administration of the DREADDs ligand JHU37160 (0.5 mg/kg, i.p.). **C.** Bar graphs showing the effects of selective activation of mPFC-LC projectors on rCPT performance metrics during the S3 probe (mPFC-LC^hM3Dq^; n=12). **D.** Bar graphs showing the effects of selective inhibition of mPFC-LC projectors on rCPT performance metrics during the S3 probe (mPFC-LC^hM4Di^; n=12). **E.** Schematic showing visual degradation of S3 stimuli to increase cognitive load during the stimulus degradation probe. **F.** Bar graphs showing impaired performance when discriminating between degraded stimuli as opposed to regular stimuli across all mPFC-LC^DREADDs^ groups (n=32). **G.** Bar graphs showing the effects of selective activation of mPFC-LC projectors on rCPT performance metrics during the SD probe (mPFC-LC^hM3Dq^; n=9). **H.** Bar graphs showing the effects of selective inhibition of mPFC-LC projectors on rCPT performance metrics during the SD probe (mPFC-LC^hM4Di^; n=8). All metrics were averaged across same-treatment sessions; error bars represent SEM. *P < 0.05, **P < 0.01.

### Inhibiting mPFC-LC projectors improves performance when cognitive load is increased by visually degrading stimuli

To further analyze the role of mPFC-LC projectors in attentional regulation, we tested the effects of manipulating their activity during conditions of increased cognitive load. Various adaptations of the rCPT have been designed to add cognitive load during task performance [25,29]. For example, we showed that increasing cognitive load by visually degrading stimuli impairs performance by reducing stimuli discrimination, and increasing FAR [26]. Given that activation of mPFC-LC projectors enhances performance by improving discrimination, we assessed the effects of manipulating mPFC-LC activity during a stimulus degradation probe (SD-probe; see Methods). Consistent with our previous findings, SD sessions produced a decrease in performance (t(31) = 9.556, p<0.0001) and response criterion (t(31) = 2.504, p=0.0178) and increased FAR (t(31) = 4.714, p<0.0001) across all mice (**Fig. 2F**). Surprisingly, J60 injections in mPFC-LC^hM3Dq^ group had no effect on performance during the SD-probe (**Fig. 2G**). Instead, J60 administration in the mPFC-LC^hM4Di^ group reduced SD induced deficits by decreasing FAR (*SAL*=0.16±0.03, *J60*=0.11±0.02; t(7) = 3.259, p=0.0139) and increasing the response criterion (*c*) (*SAL*=0.38±0.14, *J60*=0.55±0.11; t(7) = 3.702, p=0.0076) (**Fig 2H**).

### Broad activation of mPFC neurons improves performance by increasing responsiveness

To determine whether the effects of mPFC-LC activation on rCPT performance were due to specific engagement of the mPFC-LC circuit or reflected a general role of the mPFC in attentional regulation, we chemogenetically manipulated overall mPFC activity by pan-neuronally expressing either hM3Dq (mPFC^hM3Dq^), hM4Di (mPFC^hM4Di^), or mCherry (mPFC^mCherry^), and then tested the effect of manipulating mPFC activity during an S3-probe. Similar to the mPFC-LC^hM3Dq^ group, activation of DREADDs in the mPFC^hM3Dq^ group during J60 sessions significantly increased d’ (*SAL*=1.49±0.10, *J60*=1.68±0.08; t(8) = 2.331, p=0.0481; **Fig. 3C**). However, this effect was accompanied by increases in HR (*SAL*=0.60±0.04, *J60*=0.75±0.03; t(8) = 3.394, p=0.0094), FAR (*SAL*=0.14±0.02, *J60*=0.18±0.02; t(8) = 3.519, p=0.0065), total responses (*SAL*=152.20±16.98, *J60*=192.95±18.53; t(8) = 3.203, p=0.0126), as well as a decrease in the c-parameter (*SAL*=0.45±0.10, *J60*=0.12±0.09; t(8) = 3.569, p=0.0073), suggesting that improvements in rCPT performance following mPFC activation are driven by increased responsiveness and a shift toward a more liberal response strategy (**Fig. 3C**). Notably, during J60 sessions, the mPFC^hM3Dq^ group showed an increase in ITI touches (*SAL*=104.95±13.96, *J60*=141.55±18.68; t(8) = 2.302, p=0.0468), indicating heightened impulsivity (**Fig. 3C**). Interestingly, J60 administration in mPFC^hM4Di^ group also significantly increased ITI touches (*SAL*=97.40±11.47, *J60*=123.55±13.94; t(9) = 2.904, p=0.0175), but there was no change in other performance metrics (*d’: SAL*=1.34±0.13, *J60*=1.34±0.12; t(9) = 1.210, p=0.2570; *HR: SAL*=0.54±0.05, *J60*=0.55±0.06; t(9) = 0.3524, p=0.7327; *FAR: SAL*=0.12±0.01, *J60*=0.14±0.02; t(9) = 0.6047, p=0.5603; *c: SAL*=0.54±0.10, *J60*=0.51±0.13; t(9) = 0.2966, p=0.7735; *Responses: SAL*=129.55±11.02, *J60*=128.10±13.28; t(9) = 0.2370, p=0.8180)(**Fig. 3D)**. Similar to mPFC-LC^mCherry^ groups, during J60, mPFC^mCherry^ groups showed no effect on rCPT performance metrics (*d’: SAL*=1.09±0.06, *J60*=1.22±0.10; t(7) = 0.6692, p=0.5248; *HR: SAL*=0.51±0.04, *J60*=0.55±0.05; t(7) = 0.9830, p=0.3584; *FAR: SAL*=0.16±0.03, *J60*=0.15±0.02; t(7) = 0.6196, p=0.5551; *c: SAL*=0.52±0.09, *J60*=0.48±0.11; t(7) = 0.4768, p=0.6480; *Responses: SAL*=124.19±, *J60*=131.69±11.56; t(7) = 0.9564, p=0.3707)(**Fig. 3D**).

**Figure 3.**
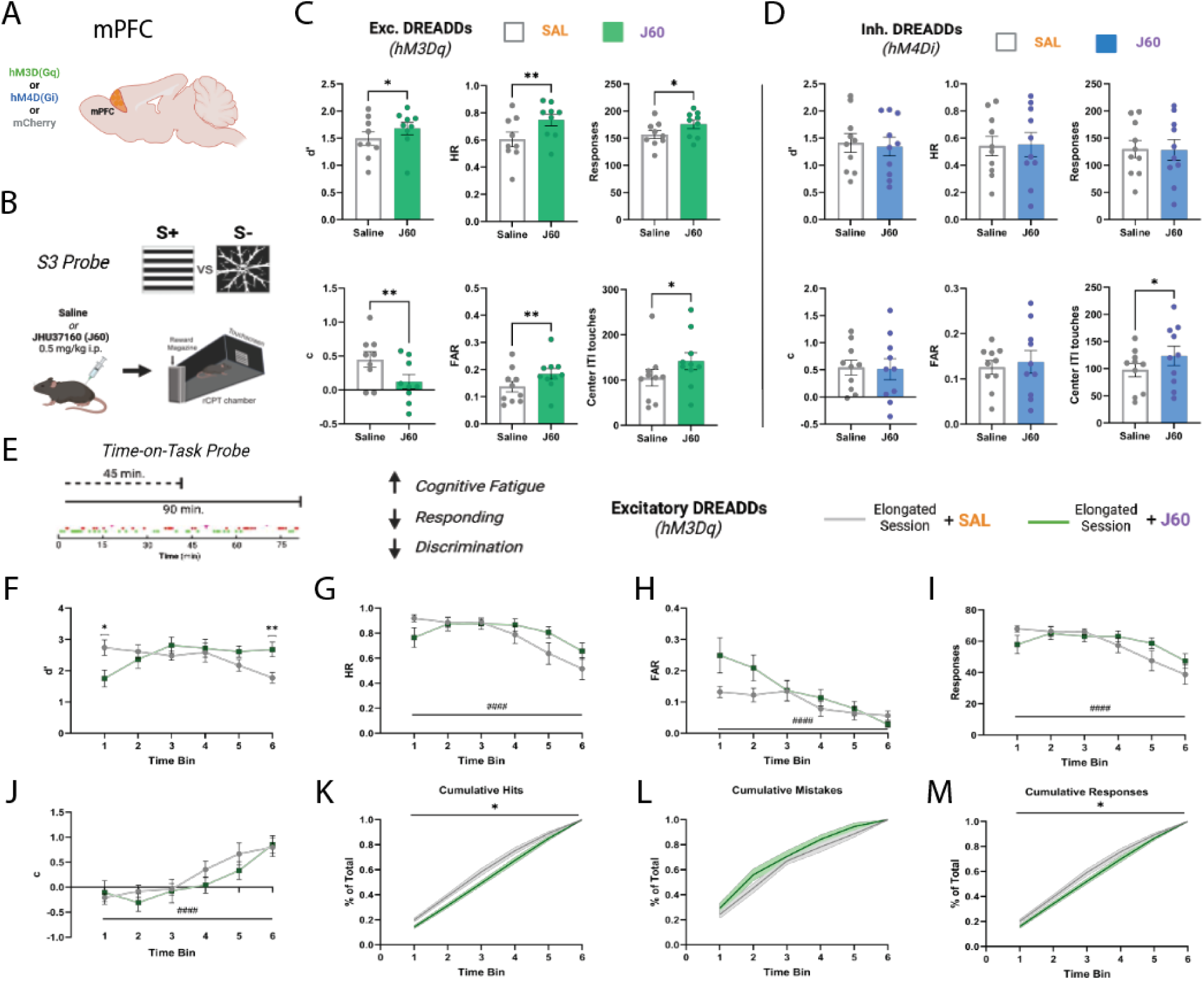
mPFC-LC inhibition reduces mistakes when cognitive load is increased through stimulus degradation. **A.** Schematic showing the expression of excitatory DREADDs (hM3Dq), inhibitory DREADDs (hM4Di) or mCherry fluorophore in the mouse mPFC. **B.** Overview of S3 experimental probe with target (S+) and non-target (S-) stimuli as well as experimental timeline for administration of DREADD ligand J60 and rCPT session. **C.** Bar graphs showing the effects of broad activation of the mPFC on rCPT performance metrics during the S3 probe (mPFC^hM3Dq^; n=10). **D.** Bar graphs showing the effects of broad inhibition of the mPFC on rCPT performance metrics during the S3 probe (mPFC^hM4Di^; n=10). All metrics were averaged across same-treatment sessions; error bars represent SEM. **E.** Schematic overview of TOT probe showing elongation of sessions from 45 to 90 minutes and a representative plot of responses over a 90 minute session. Hits are represented by points in green and false alarms are represented in red. **F-J.** Line graphs plotting rCPT performance metrics across each of 6 (15-minute) time bins during extended sessions. All metrics were averaged by time bin across same-treatment sessions; error bars represent SEM. **K-M.** Plots showing the accumulation by percent of total hits (**K**), mistakes (**L**), and responses (**M**) across time bins. All metrics were averaged by time bin across same-treatment sessions; shaded areas represent SEM. *Treatment effects:* *P < 0.05, **P < 0.01; *Time effects:* #P < 0.05, ##P < 0.01, ###P < 0.001, ####P < 0.0001.

### Broad activation of mPFC neurons reduces TOT performance decrements

Extended sessions lead to time-on-task (TOT) deficits in responsiveness and discrimination as cognitive fatigue increases and motivation decreases over time [30,31]. Given that general mPFC activation improves performance by increasing responsiveness, we hypothesized that general mPFC activation would reduce TOT deficits. To test this hypothesis, we chemogenetically manipulated mPFC activity and analyzed behavioral performance during a TOT probe (see Methods). As expected, during saline sessions, mice exhibited time-dependent declines in d′ (F (3.880, 54.32) = 6.756, p=0.0002), HR (F (2.375, 33.24) = 17.42, p<0.0001), FAR (F (2.251, 31.51) = 6.237, p=0.0040), and overall response count (F (2.482, 34.75) = 17.96, p<0.0001) across the extended session (**Fig. 3F-I**). Conversely, the *c*-parameter increased over time (F (3.073, 43.03) = 17.73, p<0.0001; **Fig. 3J**), indicating a more conservative response bias as the session progressed. During J60 sessions, mPFC^hM3Dq^ mice exhibited a significant attenuation of TOT deficits as evidenced by an increase in d’, particularly towards the final 15 min of the TOT session compared to the saline session (SALvsJ60, p=0.0077)(**Fig. 3F**). Interestingly, during the first 15 min of the session, mPFC^hM3Dq^ mice showed a decreased d’ (SAL vs J60, p=0.0159) which appears to be driven by a marginal increase in false alarms (SAL vs J60, p=0.0529). Importantly, mPFC^mCherry^ mice showed no significant differences in any metric during J60 sessions compared to saline sessions (*d’:* F (1, 12) = 0.05804, p=0.8137; *HR:* F (1,12)=0.02302, p=0.8819; *FAR:* F (1, 12) = 0.0005639, p=0.9814; *c:* F (1, 12) = 0.03218, p=0.8606; *Responses:* F(1,12)=0.008174, p=0.9295) **(Fig. S4)**.

Given that response totals can vary more drastically during elongated sessions, we analyzed the accumulation of hits, FAs, and total responses as a percentage of the session’s total within 15 min time bins to visualize their distribution over time. Broad mPFC activation shifted the distribution of hits towards later time bins (F (1, 16) = 8.306, p=0.0108; **Fig. 3J**), while FA distribution was unchanged (F (1, 16) = 2.145, p=0.1624)(**Fig. 3K**). This increase in the hits distribution towards the end of the sessions shifted overall response accumulation towards later in the session (F (1, 16) = 4.541, p=0.0489; **Fig. 3L**) and accounts for the increase d’ in the last time-bin.

### Chemogenetic manipulation of the mPFC modulates rCPT-induced activity at subcortical projection sites

Our behavioral data demonstrate that circuit-specific activation of mPFC-LC projectors produces distinct effects on attentional performance compared to broad mPFC activation. To identify potential mechanisms underlying these differences, we examined downstream targets engaged by either circuit-specific or broad mPFC chemogenetic activation during rCPT performance. Notably, we found that mPFC-LC projectors selectively innervate the LC and peri-LC region, as indicated by dense mCherry⁺ fibers in these areas (**Fig. 4B**) and robust Fos induction in LC noradrenergic (TH⁺) neurons as well as peri-LC GABAergic (VGaT⁺) and glutamatergic (VGluT2⁺) neurons (**Fig. 4C-D**), while we observed no mCherry⁺ fibers in other regions implicated in attentional control, such as the nucleus accumbens (NAc) or ventral tegmental area (VTA) (**Fig. 4B**). In contrast, mCherry expressing fibers were found in the NAc, VTA, and LC/Peri-LC region of mPFC^DREADDs^ mice (**Fig. 5A**), suggesting that these brain regions might have a role in the behavioral effects during rCPT after broad manipulation of mPFC neuron activity. Notably, broad activation of mPFC neurons induced Fos expression in the LC, peri-LC (**Fig. 5H**), and NAc (**Fig. 5D**). Interestingly, when comparing the recruitment of LC noradrenergic neurons across conditions, circuit-specific activation of mPFC-LC projectors produced a greater number of Fos⁺ LC TH⁺ neurons than broad mPFC activation (F(2, 10) = 21.06, p=0.0003, **Fig. 5I**), suggesting that targeted engagement of this pathway drives a more potent and selective activation of LC-NE neurons.

**Figure 4.**
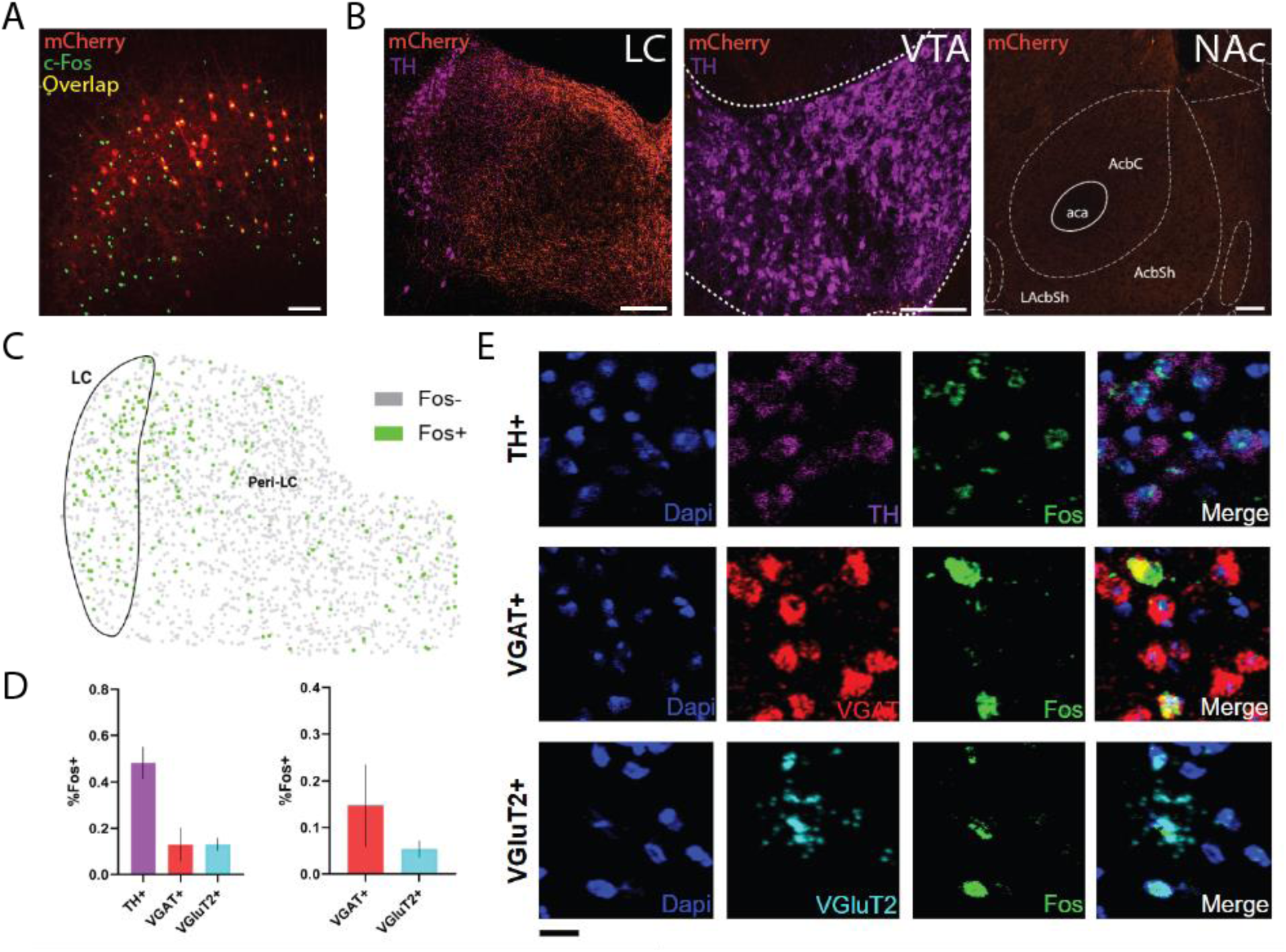
mPFC-LC projectors selectively engage a heterogeneous microcircuit in the LC and adjacent peri-LC. **A.** Image of mPFC showing expression of c-Fos (green) in mPFC-LC projectors (red) and the surrounding local microcircuit following J60 administration in mPFC-LC^hM3Dq^ mice. **B.** Representative images showing dense mCherry-expressing fibers from mPFC-LC projectors in the LC and Peri-LC. By contrast, fibers are absent from the NAc and VTA. Scale bars (white) = 200µm. **C.** Example diagram showing the distribution of *Fos*+ nuclei in the LC and peri-LC regions labeled by *in situ* hybridization following mPFC-LC activation. *Fos*+ nuclei are labeled green while *Fos*- nuclei are gray. **D.** Proportion of Fos+ nuclei by cell type in the LC and Peri-LC. Data represent an average proportion per animal and error bars represent SEM (n=3). **E.** Representative images illustrating colocalization of Fos with cell type markers for noradrenergic (TH+), GABAergic (vGAT+), and glutamatergic (vGluT2+) neurons in the LC and peri-LC region. Scale bar (black) = 10µm.

**Figure 5.**
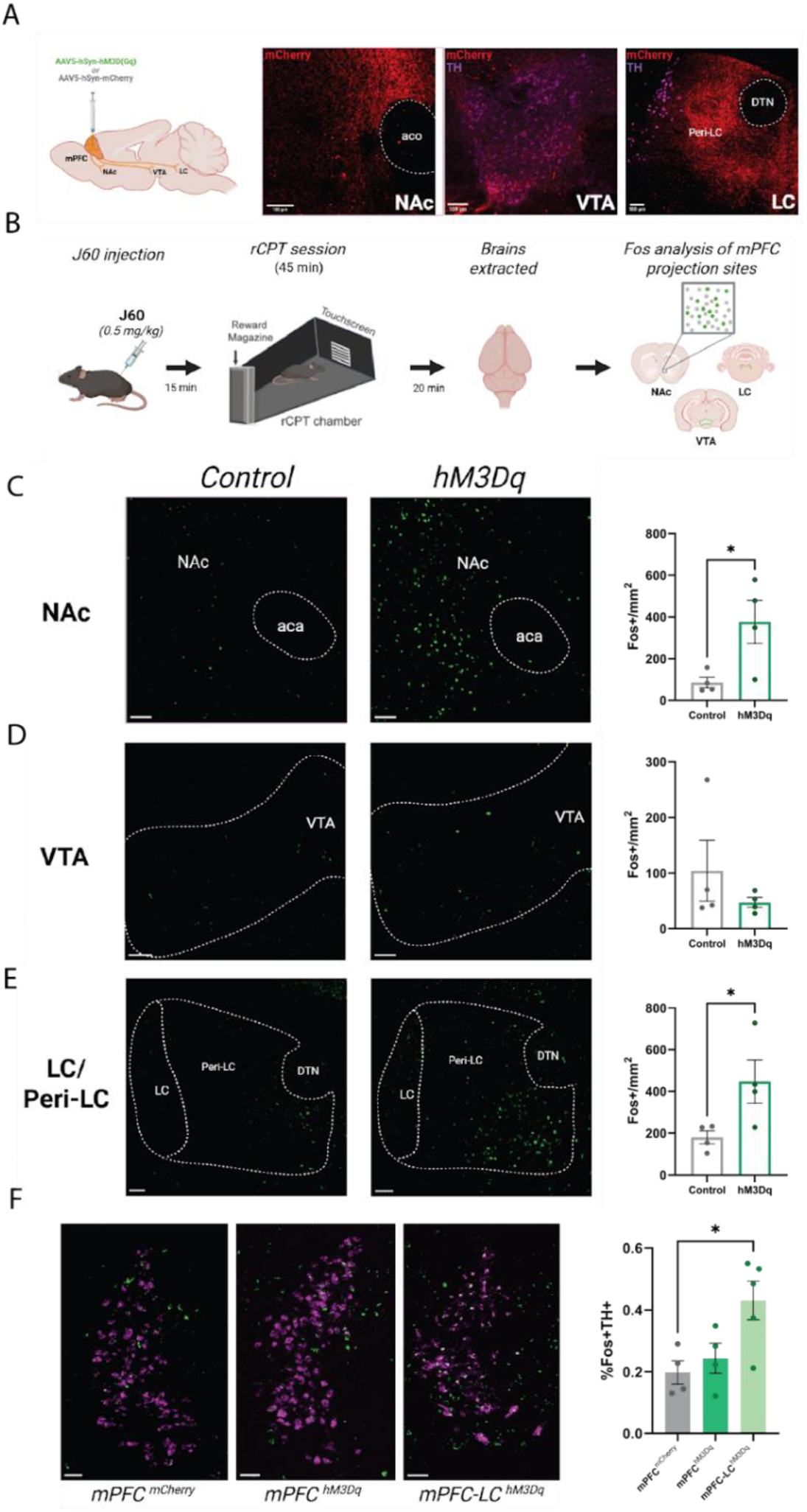
General mPFC activation modulates downstream rCPT-induced activity in attention-relevant projection sites. **A.** Schematic showing viral expression of either excitatory DREADDs (hM3Dq) or mCherry pan-neuronally in the mPFC along with representative images of mCherry-expressing fibers in the NAc, VTA, and LC. **B.** Experimental workflow for *Fos* expression-based analysis following chemogenetic manipulation during rCPT. **C-E.** Representative images of *Fos* expression following rCPT in mPFC^mCherry^ and mPFC^hM3Dq^ mice in the NAc (**C**), VTA (**D**), and LC/peri-LC region (**E**) with bar graphs showing the relative number of Fos+ nuclei per mm^2^. mPFC activation in mPFC^hM3Dq^ mice significantly increased *Fos* expression in the NAc and LC/peri-LC region (n=4 per group). Scale bars = 200µm. **F.** Representative images showing colocalization of *Fos* with *Th* across conditions and bar graph showing the relative number of TH+ LC-NE neurons expressing *Fos* in mPFC^mCherry^, mPFC^hM3Dq^, and mPFC-LC^hM3Dq^ mice. Scale bars = 100µm. Error bars represent SEM. *P < 0.05, **P < 0.01, ***P < 0.001.

## Discussion

### mPFC-LC activation improves attentional performance by enhancing discrimination during rCPT

The LC and the mPFC are implicated in attentional behavior through their roles in regulating arousal, vigilance, and norepinephrine (NE)–mediated enhancement of cognitive flexibility [32–35]. Although previous studies examined the contribution of the mPFC and NE signaling in attentional control [20,36–40], top-down regulation of attention specifically by mPFC projections to the LC has not been investigated. Here, we used chemogenetic manipulation to functionally dissect the contribution of mPFC-LC projectors to attentional performance during the rCPT. Selective activation of mPFC-LC projectors enhanced rCPT performance by increasing discrimination accuracy and reducing FAR, while broad activation of mPFC neurons enhanced performance by increasing overall responsiveness. Further, our data showing that activation of mPFC-LC projectors during the rCPT induced *Fos* expression in a large proportion of LC-NE neurons suggest that the behavioral effects of mPFC-LC activation are likely mediated by recruitment of LC-NE neurons and the resulting increase in NE tone. In contrast, *Fos* expression in regions beyond the LC following broad activation of the mPFC suggests engagement of additional downstream circuits, particularly reward circuits encompassing the NAc.

Consistent with the interpretation that LC activation and the resulting increase in NE tone underlie the behavioral effects of mPFC-LC projector activation, optogenetic stimulation of LC neurons promotes goal-directed attention [21]. Similarly, pharmacological enhancement of NE signaling via systemic administration of atomoxetine, a selective NE reuptake inhibitor, increases discrimination accuracy and reduces FAR during rCPT [29,41], an effect that parallels the effects of mPFC-LC projector activation. One of the processes by which an increase in NE tone could enhance discrimination accuracy is through modulating adaptive gain—a process through which salient or behaviorally relevant stimuli are selectively amplified to enhance task performance [42,43]. In this context, mPFC-LC projector activation may increase perceptual sensitivity to target stimuli, by tuning NE output according to task contingencies and dynamically adjusting arousal and sensory processing [20,36–38].

During regular S3 sessions, inhibiting mPFC-LC projectors did not alter performance, but trended toward increased discrimination driven by higher HR. This pattern suggests that while mPFC-LC projector activation is sufficient to enhance performance, it may not be necessary to maintain performance under familiar task conditions. Interestingly, when cognitive demand was increased by stimulus degradation, inhibiting, rather than activating mPFC-LC projectors improved rCPT performance. This paradoxical effect may reflect changes in LC NE activity and NE tone associated with task difficulty, as LC output has been suggested to scale with subjective perceptual and cognitive load [44–46]. Under degraded stimulus conditions, an increase in LC output driven by the heightened task demands may elevate NE tone to a level that optimizes performance. In this scenario, activating mPFC-LC projectors could produce little additional benefit due to a ceiling effect, whereas reducing mPFC-driven LC excitation may rebalance LC–NE activity and thereby enhance behavioral performance. All together, the data suggest that mPFC-LC projectors might function to bidirectionally tune LC dynamics to meet task needs, underscoring a more complex mechanism by which they regulate attention.

A possible mechanism by which mPFC-LC projectors fine-tune LC NE activity during different cognitive load conditions is through the recruitment of microcircuits local to the LC. Our findings showed that mPFC-LC projectors innervate both the LC and the adjacent peri-coerulear (peri-LC) region, an area that is highly interconnected with the LC and regulates LC NE activity and LC-associated behaviors [47,48]. Moreover, mPFC-LC activation engaged peri-LC GABAergic and glutamatergic neurons, suggesting that top-down cortical inputs may modulate LC output not only through direct excitation of NE neurons, but also through microcircuit-level interactions that engage the peri-LC. It is possible that engagement of these microcircuits refines LC activity by selectively recruiting certain LC-NE subpopulations while suppressing others. This interpretation aligns with recent advances in neural recording and sequencing techniques that reveal far greater heterogeneity in LC-NE neuronal identity and function than previously appreciated, with evidence that distinct LC-NE subpopulations are associated with unique cognitive processes [49–51]. This functional heterogeneity raises the possibility that LC-NE subpopulations are organized, at least in part, by their efferent targets. Single-unit recordings in both rodents and primates show that cognitive control involves the decoupling of LC–NE subgroups into different firing modes, and selective manipulation of discrete efferent modules produces separable behavioral effects [49–52].

A limitation of this study is the temporal resolution of chemogenetic manipulations, which precludes inferences about the precise timing of mPFC–LC activity during behavioral responses. We previously showed that functional connectivity between the mPFC and LC increases following false alarms in the rCPT [28], suggesting that this circuit may contribute to performance through post-error correction. Likewise, calcium-imaging identified distinct neuronal subpopulations in the mPFC that are preferentially active either during the anticipatory period preceding responses or immediately following responses [53]. It is possible that mPFC–LC projectors align with one or both of these functionally defined subpopulations.

### Broad mPFC activation enhances responsiveness in rCPT and rescues time-on-task deficits in engagement

As noted above, broad activation of mPFC neurons during the rCPT increased performance by enhancing responsiveness and shifting behavior toward a less selective response strategy. Furthermore, mPFC activation during elongated sessions attenuated TOT–related decreases in both discrimination and responsiveness. This behavioral profile may reflect an increased motivational drive to respond during the task. This interpretation aligns with neural recording and functional studies in both animals and humans demonstrating a key role for the frontal cortex in reward-driven motivation [54–56]. The mPFC sends projections to the NAc and VTA, two regions strongly implicated in motivation and reward processing [54–58]. Our data show that broad mPFC activation induced *Fos* in the NAc in addition to the LC and peri-LC, supporting the involvement of these circuits in the enhanced motivational drive underlying the increase in performance. Future studies will be needed to determine the specific contributions of the mPFC-NAc circuit in mediating the observed behavioral effects of broad mPFC activation. However, existing evidence showed that optogenetic activation of mPFC–NAc projections increase responsiveness in Pavlovian conditioning paradigms [54–56] linking this circuit with increased motivational drive.

Notably, alongside the increase in responsiveness following broad activation of mPFC neurons, we also observed an increase in ITI touches, reflecting heightened impulsivity. Interestingly, this increase in impulsivity was likewise observed after broad inhibition of the mPFC. The mPFC participates in response inhibition [59–61], and chemogenetic inhibition of the ACC [20,36–40] or mPFC-NAc projections [62] increases impulsivity. Our finding that both general mPFC activation and inhibition increased impulsivity suggests an inverse-U’ relationship of mPFC activity, in which deviations from an optimal activation state—either too high or too low—impair impulse control. Future studies will be essential to determine how this increase in impulsivity interacts with rCPT performance following broad mPFC activation. In particular, such experiments could help clarify whether the increase in responsiveness is a byproduct of heightened impulsivity and whether modifying the probability of S+ presentations alters the impact of broad mPFC manipulation on discrimination and overall rCPT performance.

**Table 1.** Viral constructs employed in the current study.

| Virus | Catalog Number (Addgene) |
| --- | --- |
| pAAV-hSyn-mCherry (AAV5) | #114472-AAV5 |
| pAAV-hSyn-hM4D(Gi)-mCherry (AAV5) | #50475-AAV5 |
| pAAV-hSyn-hM3D(Gq)-mCherry (AAV5) | #50474-AAV5 |
| pENN.AAV.hSyn.HI.eGFP-Cre.WPRE.SV40 (AAV Retrograde) | #105540-AAVrg |
| pENN.AAV.hSyn.Cre.WPRE.hGH (AAV Retrograde) | #105553-AAVrg |
| pAAV-hSyn-DIO-mCherry (AAV5) | #50459-AAV5 |
| pAAV-hSyn-hM4D(Gi)-mCherry (AAV5) | #50475-AAV5 |
| pAAV-hSyn-DIO-hM3D(Gq)-mCherry (AAV5) | #44361-AAV5 |

## Acknowledgements

We thank Haya Algrain and Stephanie Cerceo Page from the LIBD Microscopy and Spatial Biology Core for their assistance in processing and analyzing microscopy images. We also thank Suhaas Adiraju, Robert Phillips, Karen Scida, and other members of the Martinowich and Carr laboratories for helpful comments and suggestions. We also thank Aimee Ormond and Sharron Evans for assistance with animal care.

## Author Contributions

Conceptualization: KM, GVC,JMB

Formal analysis: JJR, DEO, JMB

Investigation: JJR, DEO, YL, JMB

Writing-original draft: JJR, JMB

Writing-review and editing: JJR, YL, KM, GVC, JMB

Supervision: KM, JMB, GVC

Project administration: KM, JMB, GVC

Funding acquisition: KM, GVC

## Funding

This work was supported by the National Institutes of Health award R01MH137057 (GVC,KM) and the Lieber Institute for Brain Development.

## Competing Interest

GVC is a scientific advisor for LongTermGevity, Inc. and owns stock options in the company. LongTermGevity, Inc. was not involved in the funding, design, or execution of these studies. No other authors have financial relationships with commercial interests, and the authors declare no competing interests.

---

**Figure S1.**
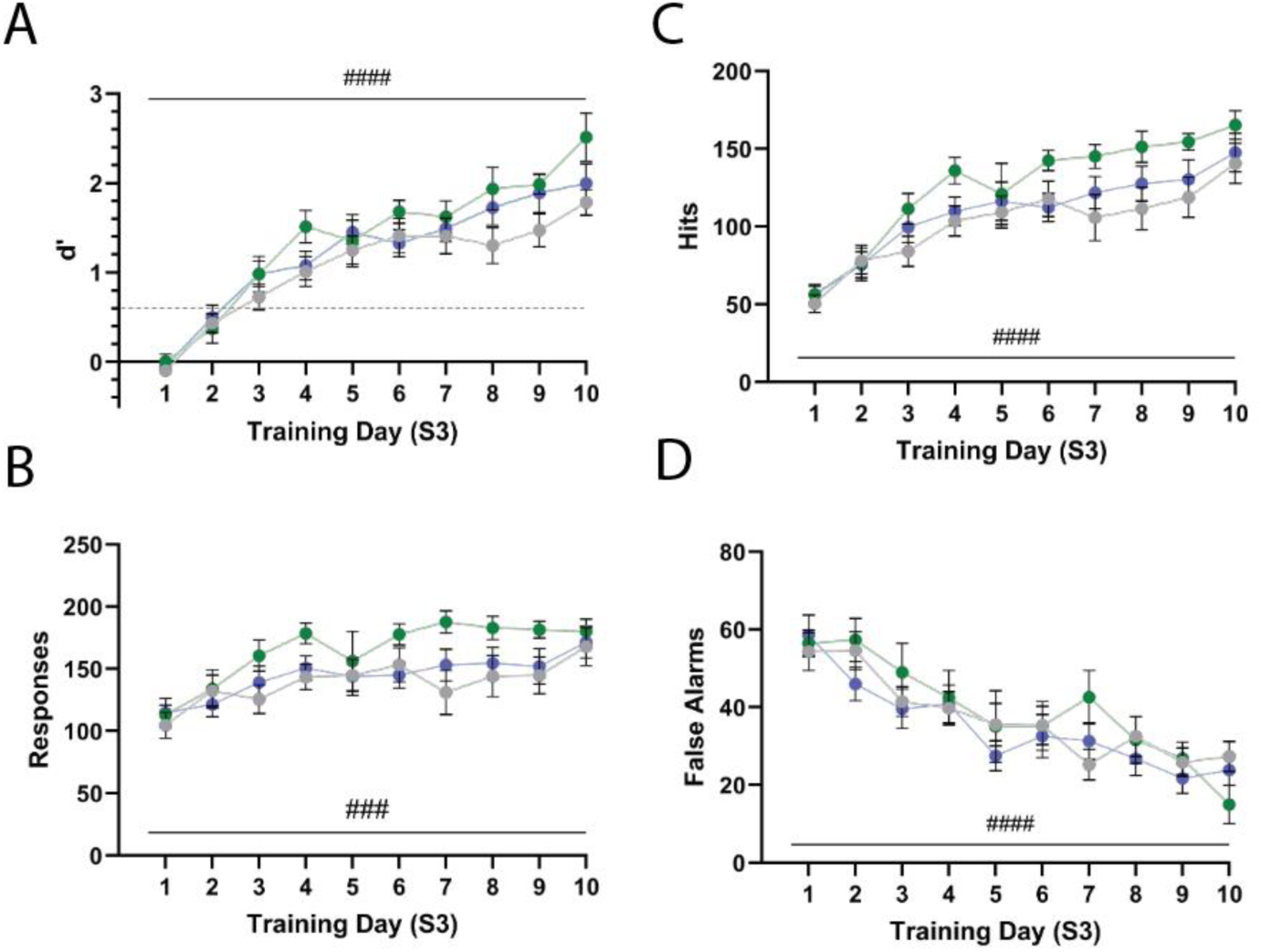
S3 training data for all experimental groups in mPFC-LC chemogenetic manipulations during rCPT. Average performance by training day for each experimental group in mPFC-LC^DREADDs^ manipulations as measured by d’ (**A**), total number of responses (**B**), number of hits (**C**), and number of false alarms (**D**)(*hM3Dq:* n=12, *hM4Di:* n=12, *mCherry:* n=8). PFC-LC^hM3Dq^ mice are represented in green, mPFC-LC^hM4Di^ mice are represented in blue, and mPFC-LC^mCherry^ mice are represented in gray. All metrics were averaged by training day across stage 3 sessions; error bars represent SEM. #P < 0.05, ##P < 0.01, ###P < 0.001, ####P < 0.0001.

**Figure S2.**
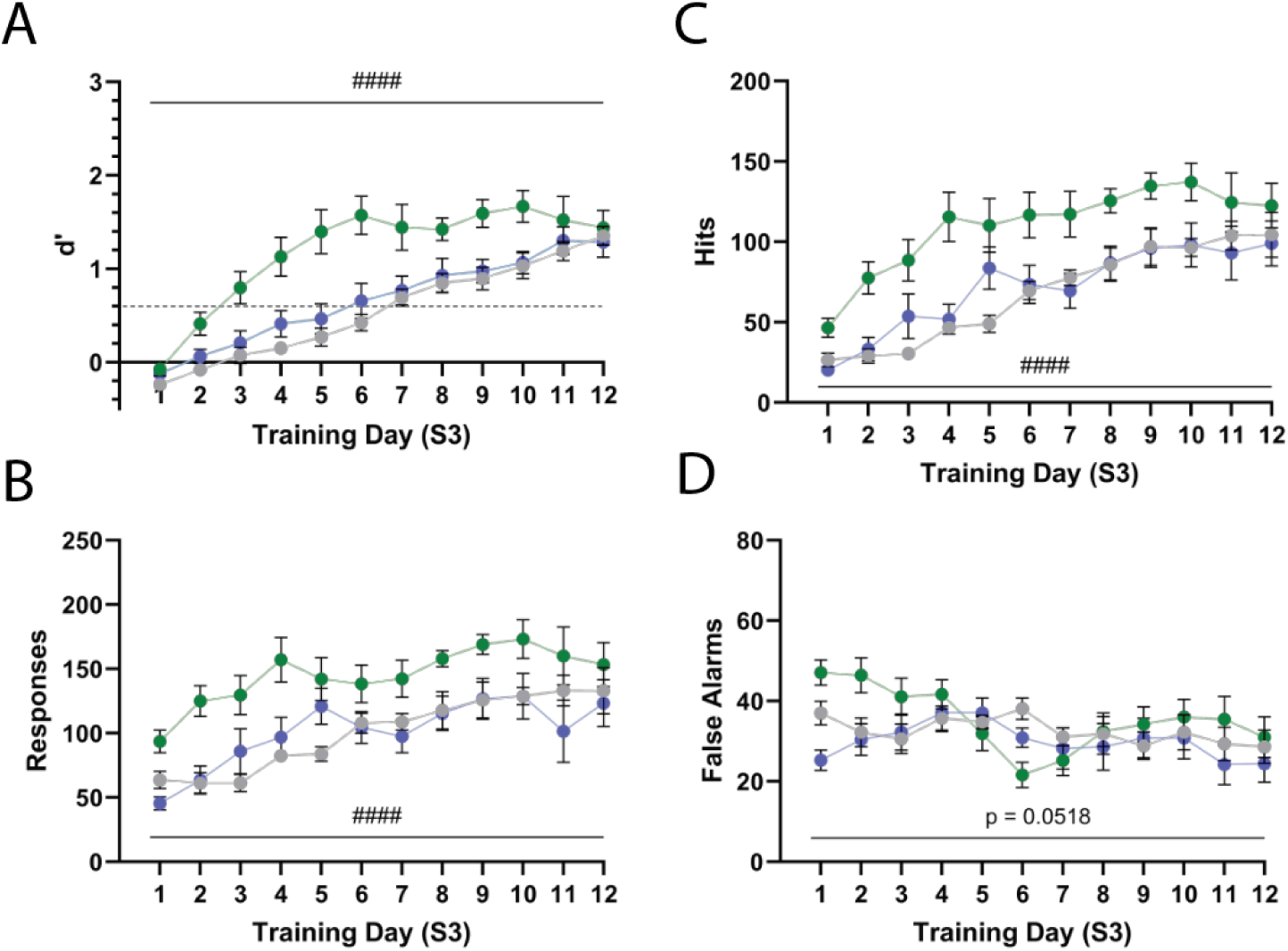
S3 training data for all experimental groups in mPFC chemogenetic manipulations during rCPT. Average performance by training day for each experimental group in mPFC^DREADDs^ manipulations as measured by d’ (**A**), total number of responses (**B**), number of hits (**C**), and number of false alarms (**D**)(*hM3Dq:* n=10, *hM4Di:* n=10, *mCherry:* n=8). mPFC^hM3Dq^ mice are represented in green, mPFC^hM4Di^ mice are represented in blue, and mPFC^mCherry^ mice are represented in gray. All metrics were averaged by training day across stage 3 sessions; error bars represent SEM. #P < 0.05, ##P < 0.01, ###P < 0.001, ####P < 0.0001.

**Figure S3.**
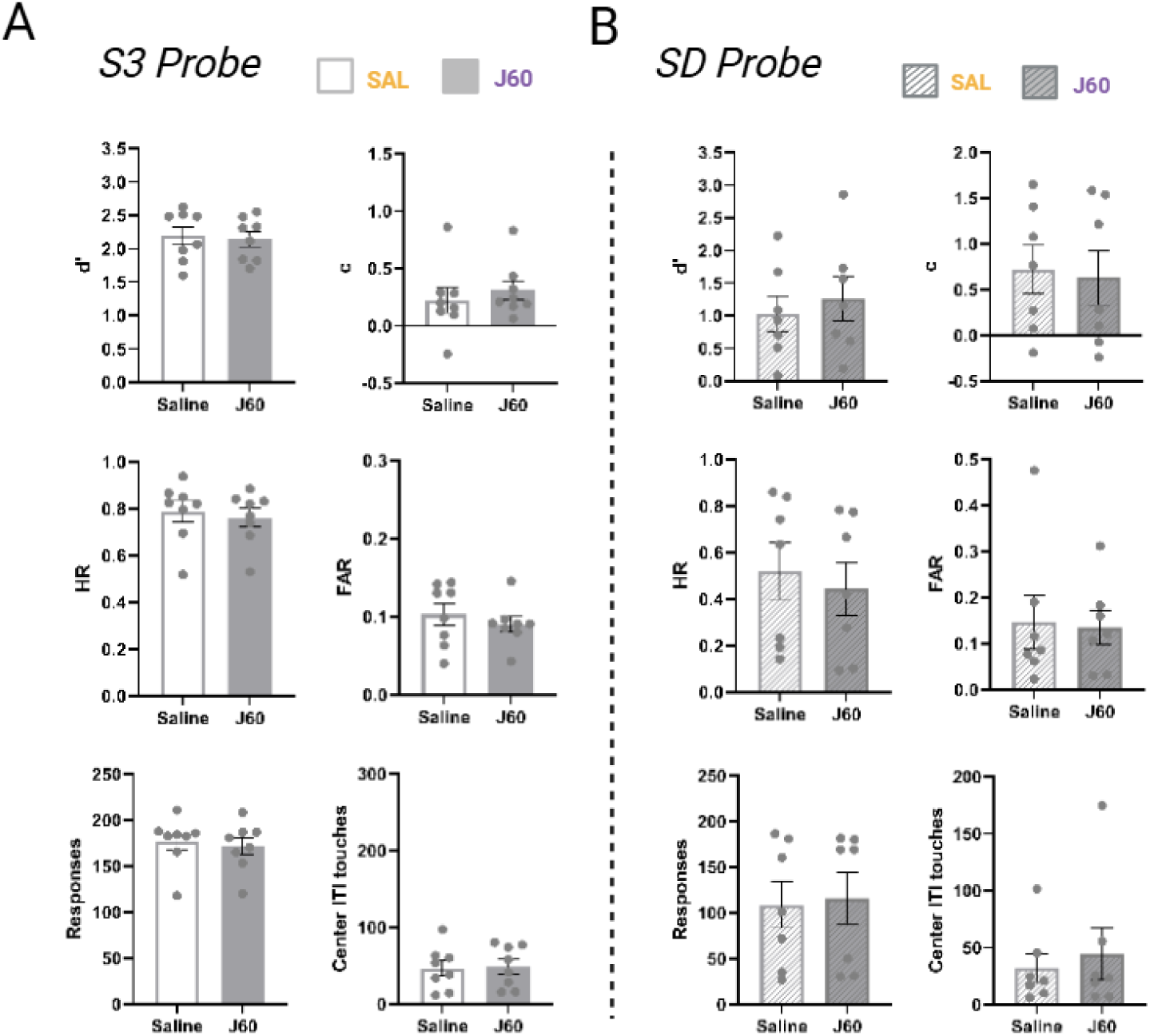
J60 has no effect on performance of mPFC-LC^mCherry^ mice in the S3 or stimulus degradation (SD) rCPT behavioral probes. **A.** Bar graphs showing comparative performance in rCPT behavioral metrics of mPFC-LC^mCherry^ mice in SAL and J60 sessions during the S3 probe (n=8). **B.** Bar graphs showing comparative performance in rCPT behavioral metrics of mPFC-LC^mCherry^ mice in SAL and J60 sessions during the stimulus degradation (SD) probe (n=7). All metrics were averaged across same-treatment sessions; error bars represent SEM. *P < 0.05, **P < 0.01.

**Figure S4.**
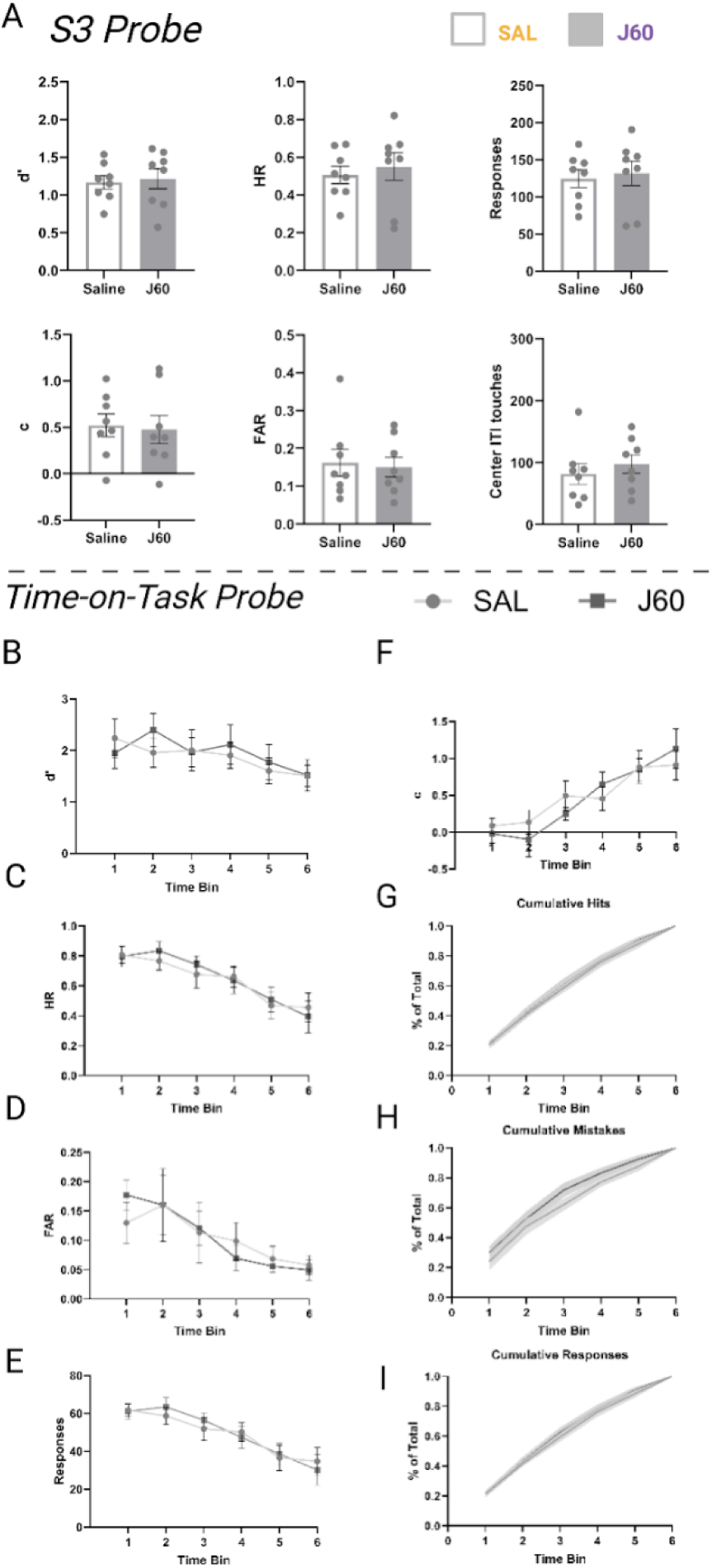
J60 has no effect on performance of mPFC^mCherry^ mice in the S3 or time-on-task (TOT) rCPT behavioral probes. **A.** Bar graphs showing comparative performance in rCPT behavioral metrics of mPFC^mCherry^ mice in SAL and J60 sessions during the S3 probe (n=8). **B-F.** Line graphs showing comparative performance in rCPT behavioral metrics of mPFC^mCherry^ mice by 15 min time bin in SAL and J60 sessions during extended sessions in the time-on-task (TOT) probe (n=8). All metrics were averaged across same-treatment sessions; error bars represent SEM. *P < 0.05, **P < 0.01.

**Figure S5.**
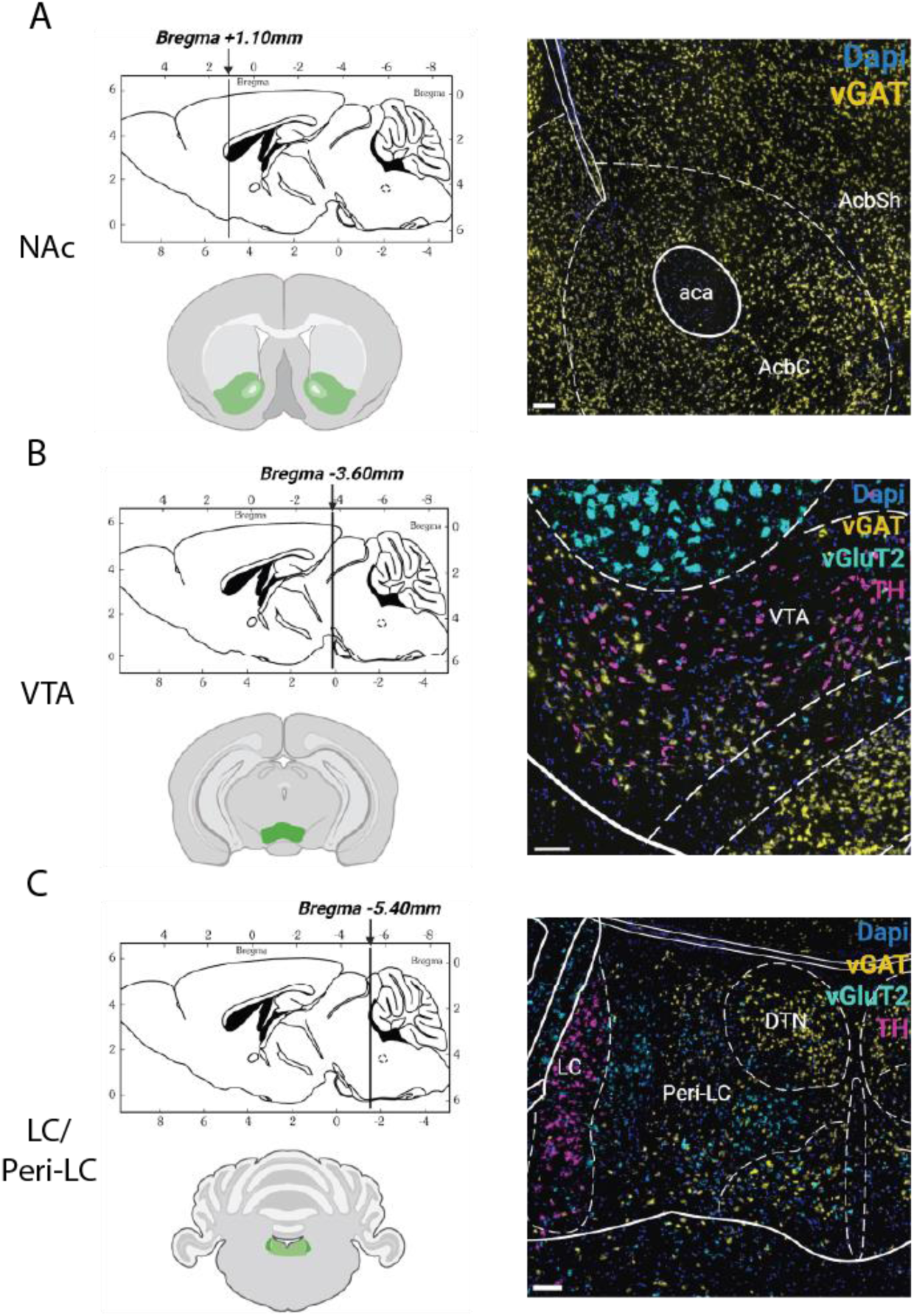
Anatomical registration of coordinates for *Fos* analysis at subcortical mPFC projection sites. Schematics and images showing the localization and alignment of slices in the NAc (**A**), VTA (**B**), and LC/Peri-LC region (**C**). Sagittal diagrams (top-left) show the coronal plane from which sections were taken. The regions analyzed are highlighted in green on schematics of coronal sections (bottom-left). Excitatory (*vGluT2*), inhibitory (*vGAT*), and catecholeminergic (*Th*) cell-type markers align with atlas-annotated topology of the respective regions. Scale bars = 200µm.

## Notes

### Summary of Updates

Updated figure 4 and figure 5

