## Supplementary figures and images for "Top-down control of sustained attention by the medial prefrontal cortex (mPFC) - locus coeruleus (LC) circuit during the rodent continuous performance test (rCPT)"

### fig S1

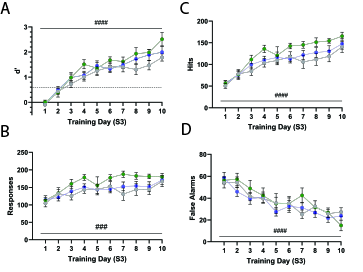

### fig S2

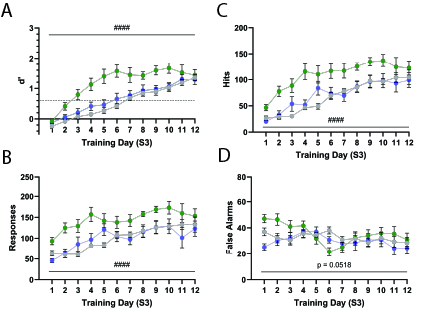

### fig S3

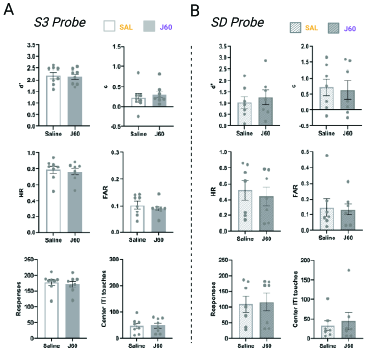

### fig S4

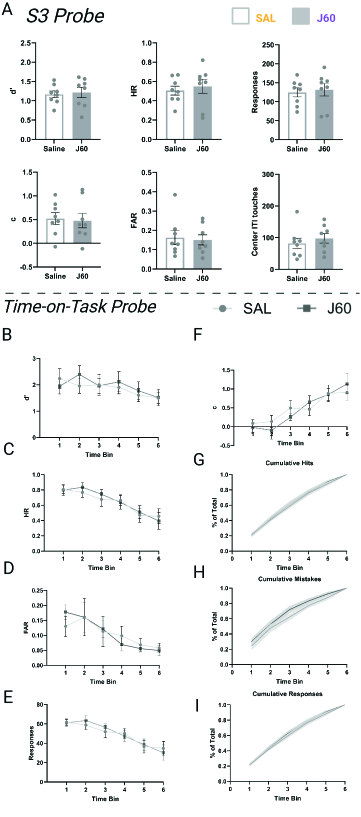

### Fig S5

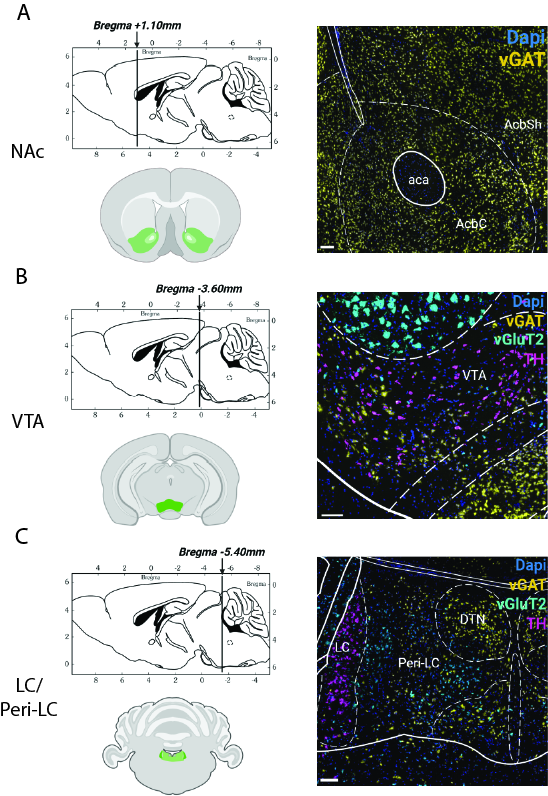
